# Does Complexation of Plasma Membrane Cholesterol with Phospholipids Determine its Transbilayer Distribution?

**DOI:** 10.1101/2025.11.13.687888

**Authors:** Theodore L. Steck, Yvonne Lange

## Abstract

The transbilayer distribution of cholesterol in the plasma membrane is unresolved in that diverse analyses have yielded contradictory results. We now propose a model based on a novel mechanism. It assumes that (a) sterols associate stoichiometrically with phospholipids and (b) these complexes are filled to capacity in the plasma membrane. The sterol in each leaflet is then the product of the abundance of the leaflet phospholipids and their sterol stoichiometries. We applied the model to the bilayer of the human erythrocyte utilizing literature values for its cholesterol content, the abundance of phospholipids in the two leaflets and their cholesterol:phospholipid stoichiometries (1:1 mole/mole in the exofacial leaflet and 1:2 mole/mole in the endofacial leaflet). The model predicts that the outer leaflet contains about two thirds of the cholesterol in this membrane; i.e., twice that in the inner leaflet. Molecular dynamics simulations have delivered similar sidedness values for membranes in which the exofacial phospholipids have a higher sterol affinity. However, our mechanism holds that sterol affinity is irrelevant if the phospholipids are replete with sterol. Our results meet the requirement that the areas of the two leaflets be about equal. The model predicts that the sterol abundance in one leaflet of a bilayer will exceed that in the other when its phospholipids are fully complexed and have a higher sterol stoichiometry.

## Introduction

The transbilayer distribution of the major lipids in the plasma membrane has been studied extensively, particularly in the human erythrocyte (Doktorova et al., 2025; Kobayashi, 2023; Lorent et al., 2020; Murate et al., 2015; Murate and Kobayashi, 2016; Pabst and Keller, 2024; Steck and Lange, 2018; Varma and Deserno, 2022). It is generally agreed that the polar lipids are asymmetrically disposed with almost all of the sphingomyelin, phosphatidylcholine and glycolipids located in the outer leaflet and phosphatidylserine, phosphatidylethanolamine and phosphatidylinositides in the cytoplasmic leaflet (Clarke et al., 2020; Lange and Steck, 2026; Murate et al., 2015; Murate and Kobayashi, 2016; Steck and Dawson, 1974; Zachowski, 1993).

Measurements of the distribution of cholesterol across the erythrocyte membrane have delivered conflicting results. These studies assign most of the sterol to one leaflet or the other or suggest a roughly even distribution (Aghaaminiha et al., 2021; Giang and Schick, 2016; Pabst and Keller, 2024; Steck and Lange, 2018). For example, opposite asymmetries were found when different quenchers were applied to the surface of cells bearing fluorescent sterol surrogates (Doktorova et al., 2025; Mondal et al., 2009; Steck and Lange, 2018; Wood et al., 2011). Consensus is also lacking for the mechanism driving the sidedness of plasma membrane cholesterol. It has been proposed that the outer leaflet holds most of the cholesterol because it is rich in saturated phospholipids with a relatively high sterol affinity (Aghaaminiha et al., 2021; Allender et al., 2019; Ingólfsson et al., 2014; Lange and Steck, 2020; Steck and Lange, 2018; Varma and Deserno, 2022). Others have argued that the sterol is pumped from the inner to the outer leaflet (Buwaneka et al., 2021; Liu et al., 2017). Another contention is that extra cholesterol compensates for a relative paucity of exofacial phospholipids so as to equalize the surface area of the two leaflets and thereby relieve interfacial tension (Doktorova et al., 2025; Varma and Deserno, 2022). It was proposed in one study that the long chain sphingomyelins in the exofacial leaflet drive cholesterol to the contralateral side (Courtney et al., 2018). Another report contended that phosphatidylserine holds the sterol in the endofacial leaflet (Maekawa and Fairn, 2015). In addition, theoretical modeling and simulations have implicated the material properties of the bilayer in determining cholesterol sidedness (Aghaaminiha et al., 2021; Allender et al., 2019; Chaisson et al., 2023; Deserno, 2024; Giang and Schick, 2016; Varma and Deserno, 2022; Yesylevskyy and Demchenko, 2012). Still other factors have been considered (Wood et al., 2011).

We now propose a novel mechanism governing the transbilayer distribution of plasma membrane cholesterol. Our hypothesis assumes that sterols form discrete associations with phospholipids. After McConnell et al., these associations are termed *complexes*. Unlike covalent and coordination complexes, they are presumably rapidly reversible stereochemical associations such as some ligands form with proteins (Anderson and McConnell, 2002; McConnell and Radhakrishnan, 2003; Radhakrishnan and McConnell, 2005; Soloviov et al., 2020). A second assumption is that the phospholipids in the two leaflets of the plasma membrane are replete with cholesterol; i.e., kept homeostatically at their stoichiometric equivalence point. In the absence of other factors, these two features set the transbilayer distribution of the sterol. Evidence supporting this hypothesis is evaluated in the Discussion.

The availability of reliable literature values for the human erythrocyte obviated the need to generate new experimental data to test the model. The application of these values leads to the prediction that about two thirds of the cholesterol in the bilayer is exofacial. An early version of this study has been published as a preprint (Steck and Lange, 2025).

## Methods

The stoichiometric complex model for bilayer sterol sidedness makes the following assumptions. (a) Cholesterol is in diffusional equilibrium within and between the liquid leaflets of the plasma membrane bilayer (Estronca et al., 2014; Steck et al., 2021). (b) Cholesterol associates stoichiometrically with the bilayer phospholipids (Lange and Steck, 2008, 2020; Lange et al., 2013; Steck et al., 2021). (c) Plasma membrane cholesterol is maintained homeostatically at the stoichiometric equivalence point of the phospholipids (Estronca et al., 2014; Lange and Steck, 2024 ; Lange et al., 2014; Steck et al., 2021). The molar cholesterol/phospholipid ratio in each leaflet is then equal to the overall stoichiometry of its complexes. (d) Consequently, the cholesterol content of each leaflet is given by the product of the abundance of its phospholipids and the stoichiometry of their complexes. (Only when the phospholipids are incompletely filled with cholesterol do their affinities affect their fractional saturation and hence the distribution of the sterol among them.)

We define C_t_, P_t_, C_o_, P_o_, C_i_ and P_i_ as the total (t) moles of cholesterol (C) and phospholipid (P) in the bilayer and in its outer (o) and inner (i) leaflets, respectively. The phospholipids in each compartment are fully complexed with cholesterol at stoichiometries n_o_ and n_i_ mole/mole_._ Thus, the resting cholesterol concentration equals its stoichiometry, denoted as C/P = C:P (mole/mole). It follows that the cholesterol in the two leaflets is given by

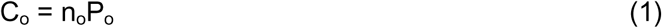

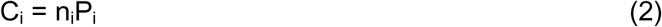

The overall bilayer cholesterol to phospholipid ratio, α, is given by:

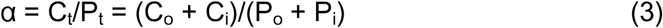

Then, substituting from Eqs. 1 and 2,

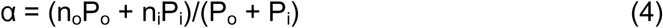

Rearranging gives

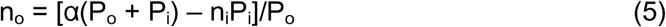

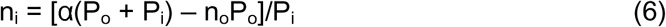

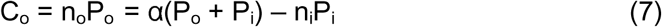

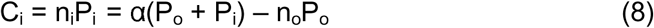

Table 1 outlines how the stoichiometric complex model predicts the concentration and transbilayer distribution of sterols in the plasma membrane. Tables 2-4 then apply the model. Rows 1 and 2 in each table give the values assumed for the amount of the phospholipid in each leaflet and their sterol stoichiometries. C_t_ is the sum of the cholesterol abundances in the two leaflets and C_t_/P_t_ is the overall bilayer concentration. The cholesterol content in each leaflet in row 3 is given by the corresponding phospholipid abundance in row 1 multiplied by its stoichiometry in row 2. Row 4 gives the cholesterol concentrations obtained by dividing row 3 by row 1. The small amounts of glycosphingolipids and minor phospholipids are ignored for want of sufficient data. We similarly disregard sterol-binding proteins; it can be estimated that, even if abundant, their stoichiometric binding of cholesterol would be negligible compared to that of the phospholipids.

**Table 1.** The stoichiometric complex model

| <u>Row</u> | <u>Parameter</u> | <u>Bilayer</u> | <u>Outer leaflet</u> | <u>Inner leaflet</u> |
| --- | --- | --- | --- | --- |
| 1 | Phospholipid, moles P | $P_t$ | $P_o$ | $P_i$ |
| 2 | C:P stoichiometry, mole/mole | $n_t$ | $n_o$ | $n_i$ |
| 3 | Cholesterol, moles C | $n_t P_t$ | $n_o P_o$ | $n_i P_i$ |
| 4 | C/P concentration, mole/mole | $C_t/P_t$ | $C_o/P_o$ | $C_i/P_i$ |

**Table 2.** The stoichiometric complex model applied to the human erythrocyte bilayer

| <u>Row</u> | <u>Parameter</u> | <u>Bilayer</u> | <u>Outer leaflet</u> | <u>Inner leaflet</u> |
| --- | --- | --- | --- | --- |
| 1 | Phospholipid, moles P | 1.00 | 0.50 | 0.50 |
| 2 | C:P stoichiometry, mole/mole | 0.75 | 1.00 | 0.50 |
| 3 | Cholesterol, moles C | 0.75 | 0.50 | 0.25 |
| 4 | C/P concentration, mole/mole | 0.75 | 1.00 | 0.50 |

**Table 3.** The stoichiometric complex model for bilayers with an alternative stoichiometry

| <u>Row</u> | <u>Parameter</u> | <u>Bilayer</u> | <u>Outer leaflet</u> | <u>Inner leaflet</u> |
| --- | --- | --- | --- | --- |
| 1 | Phospholipid, moles P | 1.00 | 0.50 | 0.50 |
| 2 | C:P stoichiometry, mole/mole | 0.70 | 0.90 | 0.50 |
| 3 | Cholesterol, moles C | 0.70 | 0.45 | 0.25 |
| 4 | C/P concentration, mole/mole | 0.70 | 0.90 | 0.50 |

**Table 4.**
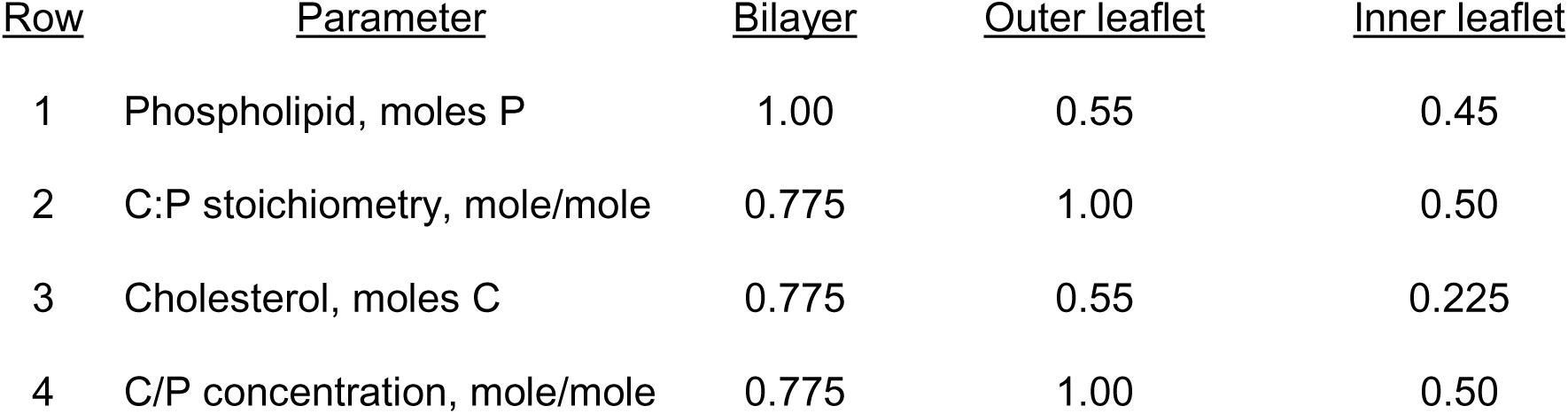
The stoichiometric complex model for bilayers with unequal P_o_ and P_i_

| <u>Row</u> | <u>Parameter</u> | <u>Bilayer</u> | <u>Outer leaflet</u> | <u>Inner leaflet</u> |
| --- | --- | --- | --- | --- |
| 1 | Phospholipid, moles P | 1.00 | 0.55 | 0.45 |
| 2 | C:P stoichiometry, mole/mole | 0.775 | 1.00 | 0.50 |
| 3 | Cholesterol, moles C | 0.775 | 0.55 | 0.225 |
| 4 | C/P concentration, mole/mole | 0.775 | 1.00 | 0.50 |

## Results

The model is tested for human erythrocyte membranes in Table 2. We express the abundance of the lipids in moles, scaled to a total bilayer phospholipid, P_t_, of one mole. In row 1, we assume that the outer leaflet is composed entirely of sphingomyelin and phosphatidylcholine (Kobayashi, 2023; Murate et al., 2015). [For a discussion of this assumption, see ref. (Lange and Steck, 2026).] These two phospholipids are assumed to contribute slightly more than half of the total phospholipid (Dodge and Phillips, 1967; Lorent et al., 2020). We therefore set the abundance of the phospholipid in the outer leaflet at P_o_ = 0.50 moles. The inner leaflet will then contain the other phospholipid species plus a few percent each of the phosphatidylcholine and sphingomyelin (Murate et al., 2015); they sum to P_i_ = 0.50 moles. Thus, in Table 2, the abundance of the phospholipid in the two leaflets is equal.

The sterol stoichiometries of dozens of the phospholipids in the human erythrocyte membrane have yet to be determined. However, representative values were assigned as follows. The outer leaflet is taken to be comprised of phosphatidylcholines and sphingomyelins; see above and ref. (Murate et al., 2015). A stoichiometry of C:P = 1.0 mole/mole was therefore assigned with confidence (Lange et al., 2013; Litz et al., 2016). This value is entered in row 2 of Table 2. The stoichiometry of the cytoplasmic leaflet phospholipids is not known and was estimated in two ways. This leaflet is enriched in unsaturated phospholipids (Lorent et al., 2020). We therefore chose dioleoylphosphatidylcholine (DOPC) as representative thereof and applied C:P = 0.5 mole/mole as its stoichiometry (Lange et al., 2013; Litz et al., 2016). In the second approach, we took C_t_/P_t_ = 0.75 mole/mole (a mole fraction of 0.43) to be the cholesterol concentration in the human red cell bilayer (Chakrabarti et al., 2017; Cooper et al., 1980; Dodge and Phillips, 1967; Doktorova et al., 2025; Lange et al., 1980). The model stipulates that this value is set by its phospholipid stoichiometry thus, C_t_:P_t_ = 0.75 mole/mole. Since it is argued that the phospholipids in the two leaflets are of equal abundance, the sterol stoichiometry of the phospholipids in the cytoplasmic leaflet can be inferred from the stoichiometry of the whole bilayer (C:P = 0.75 mole/mole) and that of its outer leaflet (C:P = 1.0 mole/mole) to be C:P = 0.5 mole/mole. This value is entered in row 2 of Table 2. (Lange et al., 2013; Litz et al., 2016; Lorent et al., 2020)

The abundance of the cholesterol in each leaflet (row 3) was estimated by multiplying the abundance of the phospholipid in each compartment (row 1) by its corresponding sterol stoichiometry (row 2; see Eqs. 1 and 2). This gives idealized estimates for the cholesterol in the outer and inner leaflets of C_o_ = 0.50 mole x 1.0 mole/mole = 0.50 moles and C_i_ = 0.50 mole x 0.50 mole/mole = 0.25 moles, respectively. These values are also those calculated using Eqs. 7 and 8. Row 4 gives the cholesterol concentrations (C/P) obtained by dividing the values in row 3 by those in row 1. Molar concentrations (i.e., C/P) can be converted to mole fractions, C/(C+P).

Row 3 in Table 2 delivers the result of interest. The predicted ratio of outer to inner leaflet cholesterol in the human erythrocyte bilayer is C_o_/C_i_ = 2. That is, the two leaflets contain 67% and 33% of the cholesterol, respectively.

## Are the areas of the two leaflets equal?

This question cannot be answered precisely because the abundance and cross-section of the dozens of the lipids in each leaflet are not known. We therefore proceeded as follows. Table 2 predicts that the outer leaflet has one third more total lipid than the inner leaflet. However, it is more condensed by roughly this factor because its phospholipid chains are more saturated. The area of the outer leaflet was calculated for two plausible compositions; one assumed 0.5 moles of dipalmitoylphosphatidylcholine (DPPC) plus 0.5 mole of cholesterol and the other assumed 0.25 moles of sphingomyelin and 0.25 moles of 1-palmitoyl-2-oleoyl phosphatidylcholine (POPC) plus 0.5 moles of cholesterol. The molecular cross-section (area of total lipid) in both cases would be ∼40 Å^2^/molecule at 30 mN/m, a surface pressure typical of plasma membranes (Aghaaminiha et al., 2021; Edholm and Nagle, 2005; Miyoshi and Kato, 2015; Stottrup et al., 2005). The relative molecular area of the exofacial leaflet was therefore taken to be approximately (0.5 moles phospholipid + 0.5 moles cholesterol) x 40 Å^2^/molecule x NA (Avogadro’s number). That is, ∼40 x NA Å^2^.

As reasoned above, we took DOPC to represent the unsaturated phospholipids in the inner leaflet and assigned it a stoichiometry of C:P = 0.5 mole/mole (Lange et al., 2013; Litz et al., 2016). This leaflet would then contain 0.25 moles cholesterol + 0.5 moles DOPC. This mixture has an overall molecular cross-section of ∼52 Å^2^/molecule at a surface pressure of 30 mN/m (Aghaaminiha et al., 2021; Dynarowicz-Łątka and Hąc-Wydro, 2004; Litz et al., 2016). The relative molecular area of the inner leaflet is then approximately (0.5 moles phospholipid + 0.25 moles cholesterol) x 52 Å^2^/molecule x NA. That is, ∼39 x NA Å^2^, nearly the same as the outer leaflet. Thus, despite the uncertainties, idealizations and approximations of these estimates, our results meet the expectation that the two bilayer leaflets have closely similar areas.

## Tests of the model

The tidy outcome in Table 2 might not be accurate or apply to other cells. We therefore tested alternative values in Tables 3 and 4. We changed the stoichiometry of the outer leaflet in row 2 of Table 3 to 0.90 mole/mole without changing the other parameters. (Such a value would obtain if the outer leaflet held a small fraction of phospholipid with a stoichiometry of C:P = 0.5 along with the predominant C:P = 1.0 species). The resulting overall membrane sterol concentration is C/P = 0.70 which is within the physiological range of C_t_/P_t_; namely, 0.70-0.80 mole/mole (mole fraction 0.41-0.44) (Chakrabarti et al., 2017; Cooper et al., 1980; Dodge and Phillips, 1967; Doktorova et al., 2025; Lange et al., 1980). In this scenario, 64% of the cholesterol is exofacial and the cholesterol asymmetry is then C_o_/C_i_ = 1.8, not far from the preferred value of 2.0. Among many other tests of our assumptions, the model predicts that outer leaflet values for C:P of less than 0.90 mole/mole yield overall C_t_/P_t_ values lower than that determined experimentally. Similarly, the assumption of outer and inner leaflet stoichiometries of C:P = 1.0 and 0.70 mole/mole leads to the prediction that C_t_/P_t_ = 0.85 mole/mole, also an unphysiologic result. Moreover, values of C:P >1.0 mole/mole have not been observed experimentally.

We also tested the possibility that the abundance of the phospholipids in the leaflets of the plasma membrane is not equal. In this case, the cholesterol concentration in the bilayer, C_t_/P_t_, is given by a general expression derived from Eq. (3),

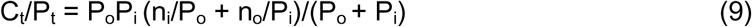

An illustration is provided in Table 4. Here, the values of P_o_ and P_i_ in row 1 were changed to 0.55 and 0.45 moles. Because P_o_ ≠ P_i_, the value for C_t_/P_t_ in row 2 is not the average of C_o_/P_o_ and C_i_/P_i_. As derived using Eq. (9), it has the value C_t_/P_t_ = 0.775 mole/mole. This prediction is well within the experimentally observed range of C_t_/P_t_ ∼0.70-0.80 mole/mole (Chakrabarti et al., 2017; Cooper et al., 1980; Dodge and Phillips, 1967; Doktorova et al., 2025; Lange et al., 1980). The ratio of outer to inner leaflet cholesterol in this case would be 2.4 with 71% of the cholesterol in the outer leaflet. Larger differences between the phospholipid content of the two leaflets did not fall within the experimental range, C_t_/P_t_ = 0.70-0.80 mole/mole.

We also tested other combinations of stoichiometries and phospholipid abundance. Some combinations predicted the observed bilayer composition. Other combinations exceeded this range in one direction or the other. We conclude that, although Table 2 presents a satisfying combination of stoichiometries and phospholipid asymmetries derived from the best literature analyses available, these values should be considered plausible, not definitive.

## Discussion

The transverse distribution of cholesterol in the plasma membrane bilayer and the mechanism by which it is established remain unresolved despite numerous studies. We approached the issue with a new mechanistic hypothesis and a simple model. The premise is that the plasma membrane bilayer phospholipids are held fully complexed with cholesterol at their stoichiometric equivalence point (Calderón and DeVries, 1997; Chakrabarti et al., 2017; Das et al., 2013; Lange et al., 1980; Lange et al., 2013; Lange et al., 2014; Litz et al., 2016; Steck et al., 2021; Symons et al., 2021; van Meer et al., 2008). The principal result is that two thirds of the cholesterol in the human red cell membrane bilayer resides in its outer leaflet (Table 2). This asymmetry agrees with that delivered by an incisive experiment based on the quenching of a fluorescent sterol surrogate in the plasma membrane of nucleated cells (Doktorova et al., 2025).

Our finding is not definitive because the sterol stoichiometries of the dozens of phospholipids in the bilayer have not been established. However, as argued in the Results, simple values from the literature provided a realistic outcome. The other major assumption, that the phospholipids in the red cell bilayer are filled to capacity, is well supported by the dramatic rise in the chemical activity (accessibility) of cholesterol incremented slightly above its resting level (Chakrabarti et al., 2017; Lange et al., 1980; Lange et al., 2013). The plasma membranes of nucleated cells similarly rest at Ct/Pt ∼0.7-0.8 mole/mole (Calderón and DeVries, 1997; Das et al., 2013; Lange et al., 2014; Steck et al., 2021; Symons et al., 2021; van Meer et al., 2008). This suggests that these membranes lipids e also are set at their stoichiometric equivalence point.

## But does cholesterol form complexes with phospholipids?

Molecular dynamics simulations typically fail to demonstrate sterol-phospholipid complexation (Aghaaminiha et al., 2021; Almeida, 2011; Chaisson et al., 2023; Varma and Deserno, 2022). However, that sterols associate with phospholipids is clear from the condensation of monolayers and bilayers (McConnell and Radhakrishnan, 2003; Regen, 2022; Róg et al., 2009). Multiple factors contribute to these associations: the shielding of the sterol molecules from water under an umbrella of phospholipid head groups; van der Waals interactions; hydrogen bonding; and polar contacts (Lange and Steck, 2008; Ohvo-Rekila et al., 2002). These forces are often characterized as a nonspecific favorable energy of interaction or excess Gibbs free energy of solvation, inferred from the partition of sterols into bilayers (Almeida, 2011; Niu and Litman, 2002; Nyholm et al., 2019; Sodt et al., 2015; Tsamaloukas et al., 2005). However, this assumption does not fit well with the data (Shaw et al., 2023).

Rather, there is diverse evidence that sterols and phospholipids do not simply mingle in the bilayer but associate stoichiometrically with stereochemical specificity. Experimental approaches include nuclear magnetic resonance spectroscopy (Darke et al., 1972), the phase behavior of binary mixtures in Langmuir monolayers (Dynarowicz-Łątka and Hąc-Wydro, 2004; Hąc-Wydro et al., 2007; Jurak, 2013; McConnell and Radhakrishnan, 2003), cryoelectron microscopy (Smothers et al., 2026), computer simulations (Jedlovszky et al., 2004; Pandit et al., 2004; Róg et al., 2009), X-ray lamellar diffraction (Hung et al., 2007), chemical cross-linking (Regen, 2022), inelastic X-ray scattering (Soloviov et al., 2020) and theoretical calculations (Radhakrishnan and McConnell, 2005). Related evidence has been reviewed (Lange and Steck, 2008; Litz et al., 2016; Quinn, 2012; Slotte, 1992; Smothers et al., 2026). Characteristic features of these complexes are described in the following paragraphs. In particular, a hallmark of sterol/phospholipid complexation is the threshold in their cholesterol-dependence isotherms illustrated by the idealized curves in Figure 1.

**Fig. 1.**
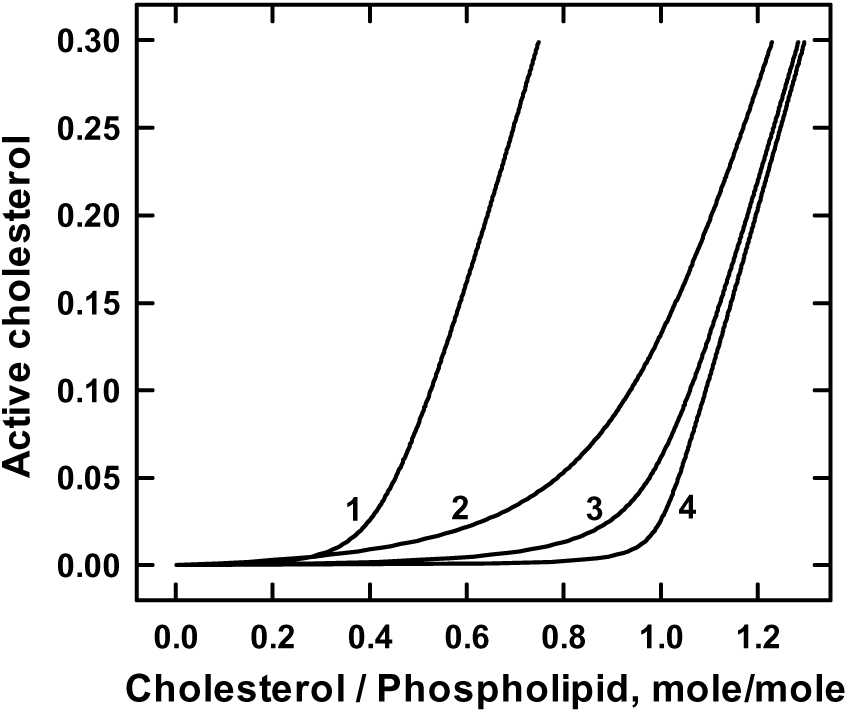
Stoichiometric associations of cholesterol and phospholipids in bilayers. Hypothetical isotherms were generated as previously described (Lange et al., 2023; Steck et al., 2024). Uncomplexed, chemically active cholesterol is plotted as a function of bilayer sterol concentration. The isotherms are taken to reflect two states of the sterol: that associated with phospholipids and the uncomplexed excess that accumulates in the bilayer at sterol concentrations exceeding the stoichiometric equivalence point. Curve 4 depicts a bilayer with a relatively high phospholipid affinity for cholesterol and a stoichiometry of C:P = 1:1 mole/mole such as observed for phosphatidylcholine (DPPC) and sphingomyelin (Lange et al., 2013). The phospholipid in curve 3 has the same 1:1 stoichiometry as curve 4 but a six-fold weaker affinity; hence, an earlier takeoff. It could represent 1-palmitoyl-2-oleoylphosphatidylcholine, POPC (Lange et al., 2013). Curve 2 illustrates competitive displacement of the sterol from phosphatidylcholine complexes by amphiphiles such as hexadecanol (Lange et al., 2009; Ratajczak et al., 2007). The stoichiometry of association in this example is the same as that in curves 3 and 4 (namely, C:P = 1:1 mole/mole) but the apparent affinity is much lower. Curve 1 has a stoichiometry of C:P = 1:2 mole/mole, that reported for DOPC (Lange et al., 2013). Its predicted threshold is therefore C/P = 0.50 mole/mole, but it takes off early due to the weak affinity of the complex.

(a) Sterol-phospholipid affinity. The relative strength of these associations varies with the fatty acyl chains and head groups of the phospholipids as well as with the structure of the sterol (Aghaaminiha et al., 2021; Lange et al., 2013; Róg et al., 2009; Rujanavech and Silbert, 1986). How affinity shapes the concentration dependence of these associations is seen in curves 3 and 4 in Figure 1. That phospholipids associate strongly with cholesterol is also manifested by their competition with cholesterol binding proteins for the sterol (Steck et al., 2024). However, this behavior is not sterol specific: phospholipids also form stoichiometric complexes with amphiphiles such as long-chain alcohols (Alanko et al., 2005; Lange et al., 2009; Ratajczak et al., 2007; Rowe et al., 1998).

(b) Sterol-phospholipid stoichiometry. Turning points in isotherms such as those pictured in Figure 1 are invariably found in the titration by sterols of the phospholipids in monolayers, liposomes and natural membranes. These thresholds signify that the complexed sterol is poorly reactive with probes and ligands whereas the super-stoichiometric excess has a high chemical activity and is increasingly available (Lange and Steck, 2008, 2020). Probes and ligands include cholesterol oxidase (Lange et al., 1980; Lange et al., 2013; Slotte, 1992), methyl-β-cyclodextrin (Ayuyan and Cohen, 2018; Litz et al., 2016; Shaw et al., 2023), bacterial toxins (Chakrabarti et al., 2017; Das et al., 2013; Lange et al., 2023) and a variety of membrane proteins (Ferrari and Tontonoz, 2025; Koh et al., 2023; Lange and Steck, 2016; Lange and Steck, 2024; Steck et al., 2024). [Note that high affinity ligands such as proteins can pull the sterol from the phospholipids; the thresholds of their binding curves therefore fall below the stoichiometric equivalence point of the sterol (Lange et al., 2023; Steck et al., 2024).] Curves 1 and 4 in Figure 1 illustrate isotherms for complexes with stoichiometries of C:P = 1:2 and 1:1, respectively.

Literature values for sterol-phospholipid stoichiometries are limited and somewhat divergent (Lange et al., 2013; Litz et al., 2016; McConnell and Radhakrishnan, 2003; Slotte, 1992). For internal consistency, we drew on a single study (Lange et al., 2013). Generally, these stoichiometries are 1:1 for saturated phospholipids and 1:2 for unsaturated phospholipids (DOPC in particular), suggesting that different phospholipids can associate with one or both faces of sterol molecules. Intermediate stoichiometries reflect mixtures of different phospholipids in the membrane. (c) Competition and displacement. Many intercalating amphiphiles increase the availability of bilayer cholesterol to probes and ligands (Alanko et al., 2005; Huang, 2009; Lange et al., 2009; Ratajczak et al., 2007; Regen, 2023). They apparently associate with the phospholipids and thereby displace the sterol from the complexes mole for mole. A comparison of curves 2 and 4 in Figure 1 illustrates such competitive displacement. The binding of cholesterol by sterol-specific binding proteins must similarly compete with the bilayer phospholipids, as seen in the sigmodicity of their sterol-dependence isotherms (Steck et al., 2024). Such competition would not occur if the sterol were simply dissolved in a phospholipid bilayer.

(d) Super-stoichiometric cholesterol has a high chemical potential. The thresholds in isotherms such as those in Figure 1 are taken to manifest the transition of the sterol in phospholipid membranes from a low to a high chemical activity as they approach their stoichiometric equivalence point (Lange and Steck, 2008, 2020; Lange et al., 2013; Litz et al., 2016; McConnell and Radhakrishnan, 2003; Radhakrishnan and McConnell, 2005). That this behavior in fact reflects a change in the chemical activity of cholesterol has been demonstrated directly (Ayuyan and Cohen, 2018; Shaw et al., 2023). Thresholds in chemical activity are characteristic of complexes but not of simple non-ideal solutions.

## Is there another mechanism for thresholds?

Do the thresholds in the concentration dependence of cholesterol isotherms reflect phase changes in the bilayer (Shaw et al., 2023)? This is unlikely, given that thresholds are observed in monolayers and vesicles containing a single phospholipid above its melting temperature (Lange et al., 2013; Litz et al., 2016; McConnell and Radhakrishnan, 2003). Nor could such thresholds be a manifestation of sterol insolubility or crystallization from a super-saturated bilayer solution. This is because these transitions occur well below the solubility limit for bilayer cholesterol; typically, C/P = 2.0 (i.e., mole fraction ∼0.67) (Huang, 2009; Litz et al., 2016). Furthermore, sterol crystals have a low chemical activity while super-stoichiometric membrane cholesterol has a high chemical activity (Ayuyan and Cohen, 2018; Shaw et al., 2023). We conclude that the thresholds observed in cholesterol isotherms lack a plausible explanation other than complexation.

## Consideration of other mechanisms driving sterol sidedness

Various theoretical treatments have explored the influence on cholesterol sidedness of stress and membrane material properties such as intrinsic and spontaneous curvature, inter-leaflet coupling and differential leaflet area (Aghaaminiha et al., 2021; Allender et al., 2019; Chaisson et al., 2023; Deserno, 2024; Giang and Schick, 2016; Varma and Deserno, 2022; Yesylevskyy and Demchenko, 2012). Our model is parsimonious and does not consider such factors. While their imposition could be additive with complexation, these forces do not appear to have much effect on relaxed human erythrocyte membranes (Himbert and Rheinstädter, 2022). Some theoretical studies have argued that the higher affinity of the exofacial phospholipids underlies the preponderance of the sterol in that leaflet (Aghaaminiha et al., 2021; Allender et al., 2019; Doktorova et al., 2025; Ingólfsson et al., 2014; Lorent et al., 2020; Nyholm et al., 2019; Steck and Lange, 2018; Varma and Deserno, 2022). There is an important difference, however, between that affinity mechanism and the stoichiometric capacity mechanism proposed here, even though the sterol asymmetry in both cases is driven by the abundance of saturated phospholipids in the outer leaflet. A differential affinity mechanism will not set the concentration of the sterol in the membrane as does the saturated complex mechanism. Since, as discussed above, plasma membrane cholesterol appears to rest at stoichiometric equivalence with the phospholipids, the complexation mechanism is favored over the affinity mechanism.

A recent publication proposed a novel mechanism driving cholesterol to the exofacial leaflet of the plasma membrane (Doktorova et al., 2025). It postulates a substantial deficit in the phospholipids in the outer leaflet with the consequent gap in its surface area being filled by excess cholesterol. This model has been critiqued (Feigenson, 2025; Lange and Steck, 2026; Skotland and Sandvig, 2026).

## Other implications of cholesterol-phospholipid complexation

If, as postulated here, the sterols and phospholipids in plasma membrane bilayers are normally close to stoichiometric equivalence, their complexes will fill both phase-separated domains (rafts) and the continuum surrounding them. Complexes would therefore contribute significantly to the material properties of the membrane: its condensation, molecular order, thickness, permeability, viscosity, compressibility, deformability, various elasticities, intrinsic curvature and lateral phase heterogeneity (Deserno, 2024; Doktorova et al., 2025; Himbert and Rheinstädter, 2022; Hofsäss et al., 2003; Khodadadi et al., 2025; Róg et al., 2009; Subczynski et al., 2017; Varma and Deserno, 2022; Vitkova and Petrov, 2013). Particular benefits conferred by the saturation of plasma membrane bilayers with sterols are a low water permeability (especially important for osmotically-vulnerable fresh water eukaryotes such as protozoa) and the imposition of a barrier against the entry of potentially deleterious amphiphiles such as xenobiotics (Khodadadi et al., 2025; Subczynski et al., 2017; Wennberg et al., 2012).

The uncomplexed fraction of plasma membrane cholesterol, although only a few percent of the total, is chemically active and performs important complementary physiological functions (Ferrari and Tontonoz, 2025; Griffiths and Wang, 2021; Lange and Steck, 2020; Lange and Steck, 2024; Sandhu et al., 2018; Steck et al., 2024; Steck and Lange, 2023; Steck et al., 2021). It is presumably the form that diffuses rapidly across the bilayer so as to equalize the chemical activity of the cholesterol in the two leaflets. In the case of the erythrocyte, equilibration of active plasma membrane cholesterol with plasma (lipo)proteins keeps the bilayer phospholipids replete at their stoichiometric equivalence point over months in circulation (Chakrabarti et al., 2017; Cooper et al., 1980; Dodge and Phillips, 1967; Doktorova et al., 2025; Estronca et al., 2014; Lange et al., 1980). The cholesterol in the plasma membrane of nucleated cells is similarly buffered by the equilibration of their active cholesterol with plasma (lipo)proteins. In addition, elaborate metabolic mechanisms maintain the plasma membrane cholesterol at stoichiometric equivalence with its phospholipids (Das et al., 2013; Lange and Steck, 2024; Steck et al., 2021). That is, super-stoichiometric plasma membrane cholesterol circulates to the cell interior and serves as a feedback signal that regulates the homeostatic elements keeping the plasma membrane phospholipids replete with cholesterol while minimizing any excess.

These considerations suggest that the stoichiometric equivalence point of plasma membrane sterol and phospholipid is a sweet spot. Less sterol undermines the functions mentioned above while a super-stoichiometric excess can be disruptive (Allen and Manning, 1996; Cooper et al., 1980; Feng et al., 2003; Kellner-Weibel et al., 2003; Subczynski et al., 2017; van Zwieten et al., 2012; Zhang et al., 2011).

## Cholesterol sidedness in other plasma membranes

Plasma membrane phospholipid composition varies among nucleated cells (Lorizate et al., 2013; Skotland and Sandvig, 2022; Symons et al., 2021). However, their sidedness is well-regulated (Norris et al., 2024) and resembles that of human red cells (Clarke et al., 2020; Doktorova et al., 2020; Kiessling et al., 2009; Kobayashi, 2023; Lorent et al., 2020; Murate et al., 2015; Murate and Kobayashi, 2016; Pabst and Keller, 2024; Zachowski, 1993). The cholesterol concentration in the plasma membranes of nucleated cells (Calderón and DeVries, 1997; Das et al., 2013; Lange et al., 2014; Steck et al., 2021; Symons et al., 2021; van Meer et al., 2008) is also similar to that in red cells, C_t_/P_t_ ∼0.7-0.8 mole/mole (Chakrabarti et al., 2017; Cooper et al., 1980; Dodge and Phillips, 1967; Doktorova et al., 2025; Lange et al., 1980). As illustrated in Tables 3 and 4, the composition and overall stoichiometry of the phospholipids in the two leaflets of plasma membrane bilayers might vary among cells but still set an asymmetrical cholesterol sidedness similar to that of the human red cell. Thus, the plasma membrane cholesterol sidedness described herein may well obtain for nucleated cells, as found in a recent study (Doktorova et al., 2025).

Intracellular membranes differ from plasma membranes in that their cholesterol levels are held well below the capacity of their phospholipids (Lange and Steck, 2024; Steck et al., 2021). The distribution of the sterol in these membranes therefore reflects the affinity of their phospholipids rather than their stoichiometry. This allows the activity of their homeostatic elements to vary with the fractional saturation of their bilayers with the sterol.

## Conclusion

If cholesterol forms complexes with bilayer phospholipids and these complexes are filled to stoichiometric equivalence in the plasma membrane then, in the absence of other influences, the cholesterol will favor the leaflet with the higher stoichiometry. This was found to be the exofacial side for the human erythrocyte membrane. Determining the phospholipid compositions, sidedness and sterol stoichiometries of a variety of other plasma membranes will test the validity and generality of this hypothesis.

## Authors contributions

TLS and YL contributed to every phase of the work from its conception to its publication.

## Acknowledgments

We are grateful to Daniel Kerr (University of Chicago), Anthony Kossiakoff (University of Chicago) and Heiko Heerklotz (University of Toronto) for their valuable comments. This work was supported by a grant for emeriti academic activities from the University of Chicago.

